# Elemental Mapping of Labeled Biological Specimens at Intermediate Energy Loss in an Energy-Filtered TEM acquired using a Direct Detection Device

**DOI:** 10.1101/2020.11.02.365940

**Authors:** Ranjan Ramachandra, Mason R. Mackey, Junru Hu, Steven T. Peltier, Nguyen-Huu Xuong, Mark H. Ellisman, Stephen R. Adams

**Author notes:** Co-senior authors and corresponding authors.

## Abstract

The multi-color or single-color EM that was developed previously, by the pseudo-colored overlay of the core-loss or high-loss EFTEM elemental map/s of the lanthanide onto the conventional image, the lanthanide chelates conjugated to diaminobenzidine being sequentially deposited as a result of selective oxidization by orthogonal photosensitizers / peroxidases. The synthesis of the new second generation lanthanide DABs, which contains 4 times more lanthanide per DAB, gives significant signal amplification and enabling collection of elemental maps at much lower energy-loss regions more favorable. Under the same experimental conditions, acquiring EFTEM elemental maps for the lanthanides at the lower energy-loss of N_4,5_ edge instead of the core-loss M_4,5_ edge, provides ~4x increase in signal-to-noise and ~2x increase in resolution. The higher signal at the N_4,5_ edge, also allows for more sophisticated technique of EFTEM spectrum Image for the acquisition of elemental maps with very high signal fidelity.

## Introduction

The Electron Microscope (EM) has gained immense popularity within the scientific community since its invention (Knoll & Ruska, 1932), partly because in addition to acquiring high resolution images, it also has the ability to simultaneously investigate the chemical/elemental composition of the sample. The elemental composition of the sample is primarily obtained either by Energy Dispersive X-ray (EDX) or Electron Energy Loss Spectroscopy (EELS), with the latter having higher sensitivity and resolution (Carter & Williams, 2009; Egerton, 1996; Goldstein, et al., 2003). The EELS technique has been applied in materials science to map elements with single atom sensitivity (Muller, 2009; Muller, et al., 2008; Varela, et al., 2005); and in biological science to detect and quantify many endogenous elements (Aronova, et al., 2008; Aronova, et al., 2009; Aronova & Leapman, 2012; Somlyo & Shuman, 1982). The EELS technique can be applied either in the Transmission Electron Microscopy (TEM) mode, generally referred to as the Energy Filtered TEM (EFTEM) (Egerton, 2012; Grogger, et al., 2005; Hofer, et al., 1997; Lozano-Perez, et al., 2009; Verbeeck, et al., 2004) or in the Scanning Transmission Electron Microscopy (STEM) mode, referred to as STEM-EELS or just EELS (Hunt, 1991;Leapman, 2017; Bosman & Keast, 2008; Goping, et al., 2003; Hunt & Williams, 1991; Kothleitner & Hofer, 2003; Leapman, 2003). Although EFTEM mode has lower sensitivity than STEM-EELS, it has at least an order of magnitude larger field of view, typically 10^5^ – 10^7^ pixels in comparison to 10^3^ – 10^5^ pixels in STEM-EELS (Aronova & Leapman, 2012; Leapman, 2017). For certain biological applications, a larger field of view is just as important as either resolution or sensitivity, as is the case for the application of multicolor electron probes to simultaneously label multiple cellular proteins/organelles in cells (Adams, et al., 2016; Pirozzi, et al., 2018; Scotuzzi, et al., 2017). In the method that we developed, the localization of the multiple targeted molecules was achieved by the sequential deposition of specific lanthanide chelates conjugated to diaminobenzidine, which were selectively oxidized by orthogonal photosensitizers / peroxidases (Adams, et al., 2016). The core-loss or high-loss (M_4,5_ edge) elemental map/maps of the lanthanides obtained by the EFTEM mode were then overlaid in pseudo-color onto a conventional electron micrograph to create the multicolor or single-color EM (Adams, et al., 2016; Boassa, et al., 2019; Sastri, et al., 2017).

The M_4,5_ core-loss edge for the lanthanides occurs at an energy loss > 800 eV. At these energy loss regions, the elemental signal is an extremely small fraction of the total incident beam (Egerton, 1996), requiring exposure times exceeding several minutes to acquire an elemental map. Such long exposures lead to poor image quality primarily due to specimen drift (Heil & Kohl, 2010), which can be somewhat offset, by taking a series of EFTEM images at shorter exposure times and then using drift correction to align them into a single image (Aoyama, et al., 2002; Heil & Kohl, 2010; Ramachandra, et al., 2014; Schaffer, et al., 2004; Terada, et al., 2001). In additional to the aforementioned drift correction strategy, we demonstrated that using a Direct Detection Devices (DDD) instead of the traditional Charge Coupled Device (CCD) to obtain EFTEM maps provided significant improvements to signal-to-noise (SNR) and spatial resolution (Ramachandra, et al., 2014). The proof of concept, of the inherent advantages of directly detecting electrons for spectroscopy, versus photo-conversion in a scintillator, was first demonstrated by Egerton, but he had to limit routine use in this mode due to the excessive beam damage (Egerton, 1984). It was only recently that beam-hardened CMOS Direct Detection Devices (DDD) able to operate continuously for months on a 300 KV TEM without significant beam damage were invented (Faruqi, et al., 2003; Faruqi, et al., 2005; Jin, et al., 2008; Milazzo, et al., 2009; Milazzo, et al., 2005; Milazzo, et al., 2010; Xuong, et al., 2007; Xuong, et al., 2004). Recently, other labs have also started to use DDD, primarily for STEM-EELS, to improve the spectral resolution and SNR (Baek, et al., 2018; Hart, et al., 2017; Maigne & Wolf, 2018).

However, in spite of these advances in EFTEM elemental mapping, we are fundamentally limited by the total intensity available at the high energy loss regions, which decreases exponentially with the increase in energy loss, in the so called Power Law form (Egerton, 1996). Alternatively, lanthanides also have a relatively strong N_4,5_ edge which occur at much lower energy loss regions, at 99 eV and 195 eV, in comparison to the M_4,5_ edge onset at 832 eV and 1588 eV for La and Lu respectively (Ahn & Krivanek, 1983). It has been suggested in the ‘EELS Atlas’, that for lanthanides the N_4,5_ edge can also be used for elemental mapping and microanalysis (Ahn & Krivanek, 1983). In continuation of our work of acquiring EFTEM elemental maps on the DDD, this paper discusses the advantages and pitfalls of obtaining elemental maps at the lower energy loss region (N_4,5_ edge) for biological samples specifically labeled with lanthanide conjugated diaminobenzidine. Technically, the energy loss region up to 50 eV is considered Low-Loss region, and the energy loss region > 50 eV is considered High-Loss region (Carter & Williams, 2009). In this paper, to make a clear distinction between the M_4,5_ and N_4,5_ edge of the Lanthanides, we will refer to the M_4,5_ edge as the high-loss (HL) region and the N_4,5_ edge as the intermediate-loss (IL) region. To facilitate these studies and future applications of color-EM, we also report the synthesis and application of novel second generation lanthanide chelates of DAB (Ln2-DAB) that deposit more metal upon DAB (photo)oxidation.

## MATERIALS and METHODS

### Materials

Reagents were obtained from Sigma-Aldrich (St. Louis, MO). Solvents and cell culture reagents were obtained from Thermo Fisher Scientific (Pittsburgh, PA) except where noted. Reactions were monitored by LC-MS (Ion Trap XCT with 1100 LC, Agilent, Santa Clara, CA) using an analytical Luna C18(2) reverse-phase column (Phenomenex, Torrance, CA), acetonitrile/H_2_O (with 0.05% v/v CF_3_CO_2_H) linear gradients, 1 mL/min flow, and ESI positive or negative ion mode. Compounds were purified by silica gel column chromatography or alternatively by preparative HPLC using the above gradients and semi-preparative Luna C18(2) columns at 3.5 ml/min.

UV–Vis absorption spectra were recorded on a Shimadzu UV-2700 (Kyoto, Japan) spectrophotometer.

### Synthesis of Ln2-DAB

#### 2,2’-Dibromo-6,6’-dinitrobenzidine (**2**)

2,2’-dinitrobenzidine (2.7 g, 10 mmol) was suspended in glacial acetic acid at 100°C with stirring, cooled to room temperature and bromine (1.13 ml, 22 mmol) in glacial acetic acid (10 ml) was added dropwise. After 30 mins at room temperature, LC-MS showed reaction was incomplete, so an additional 0.5 ml of bromine was added. After 30 min, water (100 ml) was added and the crude product collected by filtration, air-dried, and boiled in 95% EtOH (100 ml), cooled and re-filtered to give the product, **2** as an orange powder (2.34 g, 54%). ES-MS (m/z) [M]^+^, [M+ H]^+^ for C_12_H_9_Br_2_N_4_O_4_, 430.9, 432.9. Found, 431.0, 433.0.

#### Diethyl 2,2’-dinitrobenzidine-2,2’-bis-propenoate, 3

Compound 2 (1.1 g, 2.5 mmol), ethyl acrylate (2. 5 ml, 23.5 mmol), triethylamine (1 ml, 7.1 mmol). Tri(o-tolyl)phosphine (30 mg, 0.1 mmol) and palladium acetate (10 mg, 0.045 mmol) suspended in dry DMF were heated with stirring at 80°C for 3 hours and filtered hot through a glass sinter. After evaporation to dryness, the solid was suspended in 95% EtOH (50 ml) and filtered to yield the product as a brick-red solid (1.17 g, 99%). ES-MS (m/z) [M]^+^, [M+ H]^+^ for C_22_H_23_N_4_O_8_, 471.1. Found, 471.2.

#### Diethyl 2,2 ‘-dinitro-N,N’-tetra(t-butyloxycarbonyl)benzidine-2,2’-bis-propenoate, 4

Compound 3 (1.0 g, 2.1 mmol) was heated at 50°C in dry DMF (40 ml) with di-t-butyl carbonate (2.3 g, 10. 6 mmol) and DMAP (50 mg, 0.41 mmol) for 3 h. The reaction mix was evaporated and the product purified by silica gel column chromatography by eluting with 10-25% ethyl acetate-hexane to give an oil (1.69, 92%). ES-MS (m/z) [M]^+^, [M+ Na]^+^ for C_42_H_54_N_4_NaO_16_, 893.3. Found, 893.4.

#### 2,2’-Diamino-N,N’-tetra(t-butyloxycarbonyl)benzidine-2,2 ‘-bis-propanoic acid, 5

Compound 4 (1.69 g, 1.93 mmol) was hydrogenated in ethyl acetate-ethanol (1:1) with Pd/C for 6 days at rt and pressure. After filtration and evaporation to dryness, the residue was dissolved in dioxane (10 ml) and methanol (10 ml) followed by aqueous 1M-NaOH (3 ml) was then added. After overnight reaction under N_2_, the organic solvents were removed by evaporation. Water and additional NaOH was then added until all solid was dissolved. After LC-MS indicated complete saponification, glacial HOAc was added dropwise to give a precipitate that was collected by filtration after chilling in ice. The product was dried over P_2_O_5_ in vacuo overnight to give the desired product as an off-white solid (1.40 g, 96%). ES-MS (m/z) [M]^+^, [M+ H]^+^ for C_38_H_55_N_4_O_12_, 759.4. Found, 759.4. LC-MS showed 20% of the material has lost one BOC group during saponification. Tri-BOC, ES-MS (m/z) [M]^+^, [M+ H]^+^ for C_33_H_47_N_4_O_10_, 659.3. Found, 659.3.

#### 2,2’-Diamino-N,N’-tetra(t-butyloxycarbonyl)benzidine-2,2’-bis-(propanoic succinimidyl ester), 6

Compound 5 (1.40 g, ~2 mmol), N-hydroxysuccinimide (0.58 g, 5 mmol) and N-ethyl-N’-(3-dimethylaminopropyl)carbodiimide (EDC; 0. 96 g, 5 mmol) were dissolved in dry DMF and kept overnight at room temperature. The desired products (tetra and tri-Boc) were purified by prep RP-HPLC using a water-acetonitrile-0.05% trifluoroacetic acid gradient and lyophilization of the immediately frozen fractions. Yield of off-white powder, 1.14 g (60%). Tetra-Boc, ES-MS (m/z) [M]^+^, [M+ H]^+^ for C_46_H_61_N_6_O_16_, 953.4. Found, 953.4. Tri-Boc, ES-MS (m/z) [M]^+^, [M+ H]^+^ for C_41_H_53_N_6_O_14_, 953.4. Found, 953.4.

#### N-(N-2-aminoethylacetamido)-ethanediamine-N,N’,N’-triacetic acid, 7

EDTA free acid (161 mg, 0.55 mmol) was heated in dry DMSO (4 ml) under nitrogen with a heat gun until dissolved. After cooling to room temperature, EDC (106 mg, 0.55 mmol) was added with stirring. After 30 min, N-hydroxysuccinimide (63 mg, 0.55 mmol) was added followed by N-Boc ethylenediamine (174 ul, 1.1 mmol) after an additional 30 min. The reaction was kept overnight, evaporated to dryness and purified by prep RP-HPLC eluting with a gradient of water-acetonitrile-0.05% trifluoroacetic acid. Yield, 120 mg (27%) white solid. ES-MS (m/z) [M]^+^, [M+ H]^+^ for C_17_H_31_N_4_O_9_, 435.2. Found, 435.3. The BOC group was removed by dissolving the solid in trifluoroacetic acid for 30 min, evaporation and lyophilization of the resulting oil from 50% water-acetonitrile-0.05% trifluoroacetic acid.

#### Compound 8

6 (130 mg, 0.14 mmol) was added to a solution of 7 (150 mg of BOC derivative cleaved as above, 0.35 mmol) in DMSO (1 ml) with triethylamine (0.24 ml, 1.75 mmol) at room temperature. After 24h, acetic acid (0.25 ml) was added and the product separated by prep RP-HPLC eluting with a water-acetonitrile-0.05% trifluoroacetic acid gradient. Lyophilization gave a white solid, 165 mg (84%). ES-MS (m/z) [M]^+^, [M+ H]^+^ for C_62_H_95_N_12_O_24_, 1391.7. Found, 1391.8. Boc groups were removed by treating with TFA for 15 min at rt followed by immediate evaporation. The desired product was purified by prep RP-HPLC eluting with a water-acetonitrile-0.05% trifluoroacetic acid gradient. Lyophilization gave a white solid, 80 mg. ES-MS (m/z) [M]^+^, [M+ H]^+^ for C_42_H_63_N_12_O_16_, 991.4. Found, 992.0. Titration with solutions of CeCl_3_ using arsenazo III pH 7.2 as endpoint indicator in 100mM MOPS buffer gave 60% purity by weight indicating 6 TFA molecules are present in the resulting product.

#### Ln2-DAB, 9

To a 2 mg/ml (1.2 mM) solution of 8 in 0.1M sodium cacodylate pH 7.4, 25 ul LnCl_3_ (where Ln= La, Ce, Pr, Nd; 100 mM solutions in water, except Nd was a 100mM solution in 0.1N HCl) were added per ml and the pH adjusted to pH 7.4 with 1M-NaOH.

### Sample Preparation

#### Mitochondrial matrix-APEX2 labeled with Ce2-DAB (second generation lanthanide DAB) and DAB

HEK293T cells were cultured on imaging plates containing poly-d-lysine coated glass bottom No. 0 coverslips (P35GC-0-14C, MatTek Corporation). Cells were transiently transfected with Mitochondrial matrix-APEX2 fusion using Lipofectamine 3000 (Life Technologies). APEX2 was fused to C-terminal fusion of mito matrix (Martell, et al., 2012). After 16 hours transfection, cells were fixed with 2% glutaraldehyde (18426, Ted Pella Incorporated) in 0.1M sodium cacodylate buffer, pH 7.4 (18851, Ted Pella Incorporated) containing 1 mM CaCl_2_ for 5 minutes at 37°C and then on ice for 55 minutes. Fixative was removed and cells were rinsed with 0.1M sodium cacodylate buffer pH 7.4 (5X1min) on ice. On a set of plates an enzymatic reaction with 1.2 mM Ce2-DAB with 5 mM H_2_O_2_ (from 30% stock) in 0.1M sodium cacodylate buffer, pH 7.4 containing 1mM CaCl_2_ buffer solution at pH 7.4 for 5 minutes was completed. On a separate control plate of transfected cells was an enzymatic reaction of 2.5 mM DAB with 5 mM H_2_O_2_ in 0.1M sodium cacodylate buffer, pH 7.4 containing 1 mM CaCl_2_ buffer solution at pH 7.4 for 5 minutes. After reactions, all plates of cells were rinsed with 0.1M sodium cacodylate buffer pH 7.4 (5X1min) on ice and then were posted fixed with 0.100% ruthenium tetroxide (20700-05, Electron Microscopy Sciences) containing 2 mM CaCl_2_ and in 0.1M sodium cacodylate buffer, pH 7.4 for 20 minutes.

#### MiniSOG-H2B labeled with La2-DAB (second generation lanthanide DAB)

HEK293T cells were cultured on imaging plates containing poly-d-lysine coated glass bottom No. 0 coverslips (P35GC-0-14C, MatTek Corporation). Cells were transiently transfected with miniSOG-H2B fusion using Lipofectamine 3000 (Life Technologies). MiniSOG was fused to N-terminus of H2B. After 16 hours of transfection, cells were fixed with 2% glutaraldehyde (18426, Ted Pella Incorporated) in 0.1M sodium cacodylate buffer, pH 7.4 (18851, Ted Pella Incorporated) containing 1 mM CaCl_2_ for 5 minutes at 37°C and then on ice for 55 minutes. Fixative was removed, cells were rinsed with 0.1M sodium cacodylate buffer pH 7.4 (5X1min) on ice and treated for 30 min in blocking buffer consisting of 50 mM glycine, 10 mM KCN, and 5 mM aminotriazole to reduce nonspecific back-ground reaction of diaminobenzidine (DAB). Pre-equilibration with second generation La2-DAB was carried out in 0.1M sodium cacodylate buffer, pH 7.4 containing 1 mM CaCl_2_ buffer solution at pH 7.4 that was added to the plates for 30 minutes following filtration with a 0.22 μm Millex 33mm PES sterile filter (SLGSR33RS, Sigma-Aldrich) at room temperature. Photooxidation was performed by blowing medical grade oxygen over the La2-DAB solution and cells were illuminated using a standard FITC filter set (EX470/40, DM510, BA520) with intense light from a 150W xenon lamp. Illumination was stopped as soon as a light brown precipitate occurred after 8-10 minutes. After reaction, the La2-DAB solution was removed and cells were rinsed with 0.1M sodium cacodylate buffer pH 7.4 (5X1min) on ice and then were posted fixed with 1% osmium tetroxide (19150, Electron Microscopy Sciences) containing 0.8% potassium ferrocyanide, 2mM CaCl_2_ and in 0.1M sodium cacodylate buffer, pH 7.4 for 30 minutes.

#### DNA labeled with EdU and clicked with Fe-TAML-azide for oxidation of Nd-DAB2 (first generation lanthanide DAB)

Small epithelial airway cells (SEAC) were cultured on imaging plates containing poly-d-lysine coated glass bottom No. 0 coverslips (P35GC-0-14C, MatTek Corporation). SEAC were incubated with EdU (1149-100, Click Chemistry Tools) the night before for 12 hours. The cells were fixed with 2% glutaraldehyde (18426, Ted Pella Incorporated) in 0.1M sodium cacodylate buffer, pH 7.4 (18851, Ted Pella Incorporated) containing 1mM CaCl_2_ for 5 minutes at 37°C and then on ice for 55 minutes. Fixative was removed and cells were washed with 0.1M sodium cacodylate buffer pH 7.4 (5X2min) on ice, with 1X PBS (2X2min) at room temperature and rinsed 2 times quickly with 1%BSA-1XPBS by using a 0.22 μm Millex 33mm PES sterile filter (SLGSR33RS, Sigma-Aldrich) at room temperature. Cells were click reacted with 1 ml solution containing and added in the order listed, 900 μl click buffer (50 mM HEPES pH 7.6 100 mM NaCl, 0.1% saponin), 10 μl CuSO4 (100 mM), 1 μl Fe-TAML-azide (14 mM stock, giving 28 μM, Mackay et al, manuscript in preparation) and 50 μl of freshly made sodium ascorbate (100 mM) and kept covered at room temperature for 60 minutes with protection from light. During the halfway point of the click reaction, an additional 50 μl of freshly made sodium ascorbate solution (100 mM) was added to the incubation solution. Cells were then washed with filtered 1%BSA-1XPBS (2X2min) at room temperature, 1X PBS (2X2min) at room temperature, 50 mM Bicine-100mM NaCl buffer at pH 8.3 (2X2min) and reacted with 2.5 mM Nd2-DAB in 50 mM Bicine-100mM NaCl buffer solution with 40 mM H_2_O_2_ (from 30%) at pH 8.3 for 15 minutes. Cells were then washed with filtered 1%BSA-1XPBS (2X1min) at room temperature, 1X PBS (2X1min) at room temperature, 50 mM Bicine-100mM NaCl buffer at pH 8.3 (2X2min), rinsed with 0.1M sodium cacodylate buffer pH 7.4 (5X1min) on ice and then were posted fixed with 1% osmium tetroxide (19150, Electron Microscopy Sciences) containing 0.8% potassium ferrocyanide, 2mM CaCl_2_ and in 0.1M sodium cacodylate buffer, pH 7.4 for 30 minutes.

#### Specimen processing post secondary fixation

Post fixative was removed from cells and were rinsed with 0.1M sodium cacodylate buffer pH 7.4 (5X1min) on ice. Cells were washed with ddH2O at room temperature (5X1min) followed by an ice-cold graded dehydration ethanol series of 20%, 50%, 70%, 90%, 100% (anhydrous) for one minute each and 2X100% (anhydrous) at room temperature for 1 minute each. Cells were infiltrated with one part Durcupan ACM epoxy resin (44610, Sigma-Aldrich) to one part anhydrous ethanol for 30 minutes, 3 times with 100% Durcupan resin for 1 hours each, a final change of Durcupan resin and immediately placed in a vacuum oven at 60°C for 48 hours to harden. Cells were identified, cut out by jewel saw and mounted on dummy blocks with Krazy glue. Coverslips were removed and 100 nm thick specimen sections were created with a Leica Ultracut UCT ultramicrotome and Diatome Ultra 45° 4mm wet diamond knife. Sections were picked up with 50 mesh gilder copper grids (G50, Ted Pella, Inc) and carbon coated on both sides with a Cressington 208 Carbon Coater for 15 second at 3.4 volts.

### Electron Microscopy

Electron microscopy was performed with a JEOL JEM-3200EF transmission electron microscope, equipped with a LaB6 source operating at 200 KV. The microscope is fitted with an in-column Omega filter after the intermediate lenses and before the projector lenses. The Cs and Cc aberration coefficients of the objective lens are 3.2 and 3.0 mm respectively. Conventional TEM and EFTEM images were collected using a condenser aperture of size 120 μm, an objective aperture of size 30 μm and entrance aperture of size 120 μm was used. Additionally, a selective area aperture of size 50 μm was used for acquisition of electron energy-loss spectra. The spectrometer energy resolution is 2 eV, measured as the FWHM of the zero-loss peak. Before the acquisition of elemental maps, the sample was pre-irradiated with a low beam dose of ~ 3.5 × 10^4^ e-/nm^2^ for about 20 mins to stabilize the sample and to reduce contamination (Egerton, et al., 2004).

The conventional TEM images and electron energy loss spectra were acquired on an Ultrascan 4000 CCD from Gatan (Pleasanton, CA). All the elemental maps, irrespective of whether it is a high-loss or an intermediate-loss map, were acquired on a DE-12 camera, which is a DDD from Direct Electron LP (San Diego, CA). The maps presented in this paper were acquired either by the traditional 3-window method (Berger & Kohl, 1993; Egerton, 1996) or by the EFTEM Spectrum Imaging (EFTEM SI) technique (Kortje, 1994; Lavergne, et al., 1992; Schaffer, et al., 2006; Watanabe & Allen, 2012). For the 3-window method, to mitigate the effects of sample drift, instead of acquiring a single Pre-edge 1, Pre-edge 2 and Post-edge image for long exposures, a series of images of shorter durations for each of the pre-edge and the post-edge were acquired. These images were drift corrected and merged to form a single image pre-edge or post-edge image as the case may be. Also, instead of sequentially collecting the entire series of images at one energy window before proceeding to the next energy window, we acquired a set of images successively through all the energy windows and then acquire the next set through all the energy windows and so on. This interlaced acquisition reduces detrimental effects due to sample shrinkage/warping and high-tension instabilities over time, has been previously described in more detail (Adams, et al., 2016). The acquisition of both the 3-window method and EFTEM SI was accomplished by writing a macro in Serial EM for the control DE-12 detector (Mastronarde, 2005). The high-loss EFTEM images were aligned using the Template Matching plug-in in ImageJ (Tseng, et al., 2011) and the intermediate-loss EFTEM images were aligned using the filters and alignment routines of Digital Micrograph (Gatan, Inc). The elemental maps for the 3-window method was computed using the EFTEM-TomoJ plug-in of ImageJ (Messaoudi, et al., 2013) and the elemental maps for the EFTEM SI was computed using the Digital Micrograph (Gatan, Inc). The EFTEM images were dark current subtracted only and not gain normalized to avoid the fixed pattern noise arising due to uncertainties in flatfielding (Ramachandra, et al., 2014). All the elemental maps presented in this paper, were acquired at the full resolution of the DE-12 detector with out any binning applied to them. The SNR of the elemental map was obtained by dividing the signal by the standard deviation of the background intensity (He & Zhou, 2008; Waters, 2009). The signal was calculated by subtracting the mean intensity of regions that contained the lanthanide from the mean intensity of regions that represented the background. Please see the supplementary section in this paper, for the detailed TEM and spectrometer acquisition parameters for each of the datasets presented.

## RESULTS and DISCUSSION

The method for multi-color or single-color EM that we developed depends upon the sequential oxidative deposition of Ln-DAB2 (Scheme1), consisting of a single lanthanide ion, Ln^3+^ bound to the chelate (DTPA) conjugated to one of the amino groups of two diaminobenzidine molecules (Adams, et al., 2016) by two of its carboxylate groups. To deposit more lanthanide per oxidized DAB, we synthesized Ln-DAB2 (Scheme 1), by a multi-step synthesis from 3,3’-dinitrobenzidine by bromination to the 5,5’-dihalo derivative, followed by Heck reaction with ethyl acrylate. BOC-protection of the amino groups was necessary to avoid their intramolecular cyclization with the ortho acrylic esters. Catalytic hydrogenation of acrylic and nitro groups, and saponification of the ethyl ester gave the resulting diacid that was coupled to N-aminoethyl-EDTA via an intermediate bis-NHS ester, to give the BOC-protected DAB derivative conjugated to two EDTA chelates. Acid deprotection and titration with Ce^3+^ or Ln^3+^ gave the final product, Ln2-DAB, in which 2 lanthanide ions are bound per DAB and would therefore deposit four-fold more Ln^3+^ per oxidized DAB than our first generation chelate, Ln-DAB2.

HEK293T tissue culture cells transiently transfected with the genetically-encoded peroxidase, APEX2 were fixed with 2 % glutaraldehyde and incubated with Ce2-DAB and H_2_O_2_ for 5 minutes until a faint darkening of the cells was visible by bright field microscopy. Followed by post-fixation with RuO4, and then dehydrated, infiltrated, embedded and sectioned for TEM (Hayat, 1981). Previously, the EFTEM elemental maps for the lanthanide/s were obtained at their high-loss M_4,5_ edge, which were then overlaid in pseudo-color onto a conventional electron micrograph to create the so-called Color EM (see figure 1)(Adams, et al., 2016).

**Figure 1.**
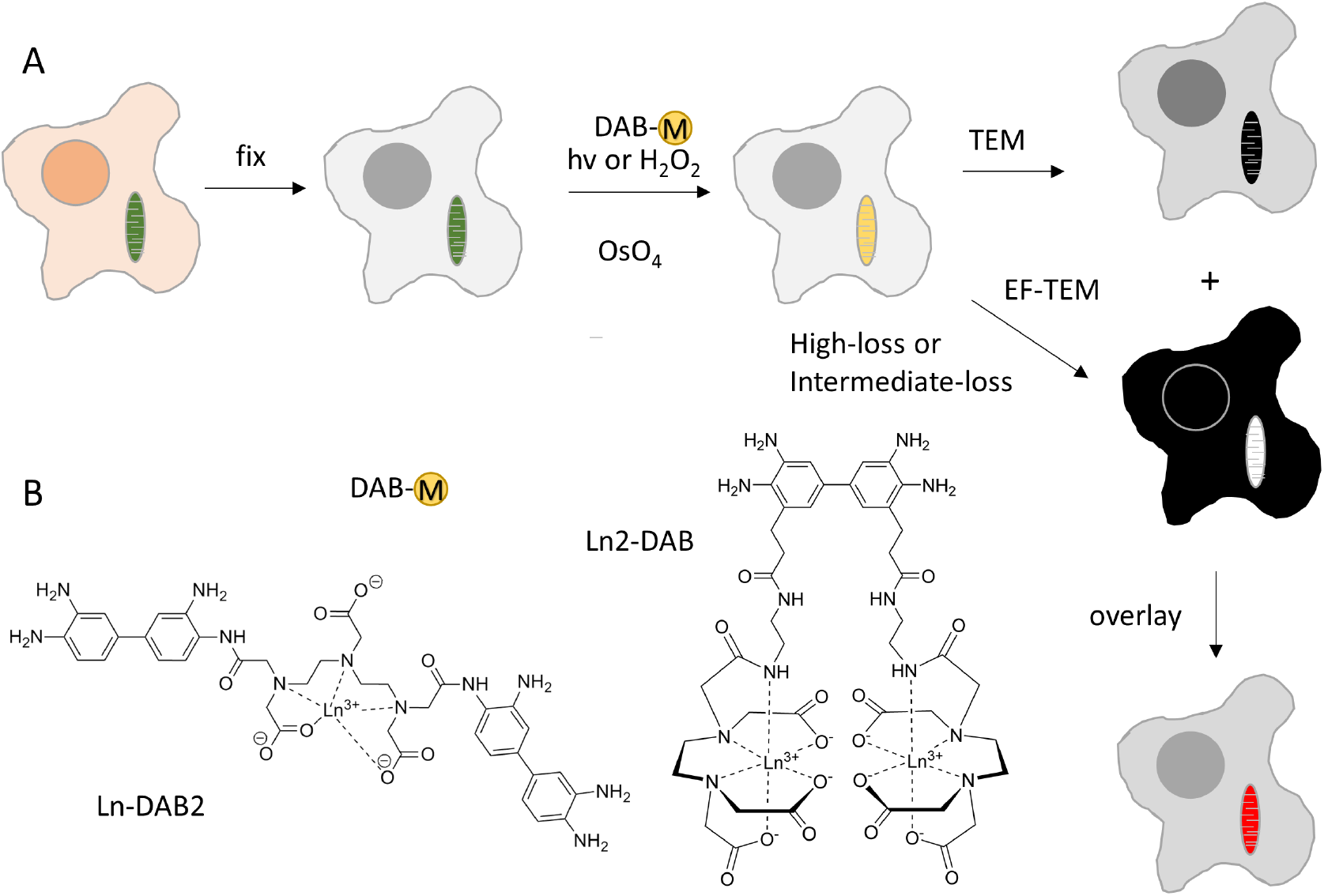
**a)** Scheme of process for obtaining color-EM images of cells. Cells containing mitochondria labeled with a photosensitizer or peroxidase are fixed, incubated with a metal chelate of DAB, DAB-M and either irradiated or incubated with H_2_O_2_ respectively. Following osmification, embedding and sectioning, corresponding TEM and EF-TEM images are collected at element-distinctive high loss and intermediate loss energies. A pseudocolor overlay of these electron energy loss images on the osmium TEM yields a color-EM image of the cells. **b)** Chemical structures of DAB-lanthanide metal chelates; first generation, Ln-DAB2 and second generation, Ln2-DAB.

The main drawback of obtaining elemental maps at the M_4,5_ high-loss region for the lanthanides, is that the total intensity is very low, and requires long exposures at relatively high dose and dose rates (Ramachandra, et al., 2014). The background intensity I varies as a function of Energy Loss E, in the so called power-law form I = AE^-r^, with values for parameters A and r being dependent on the experimental conditions (Egerton, 1996). The overall signal intensity drops exponentially with energy loss, reaching negligible levels for energy loss above 2000 eV, and which is the practical limit of the technique (Carter & Williams, 2009). For example, it has been shown that only 5 × 10^-5^ fraction of the total incident electron dose contributed to the oxygen K edge (onset at 532 eV) for a sample of 40 nm amorphous ice (Maigne & Wolf, 2018).

It has been suggested that for lanthanides, the N_4,5_ edge occurring at much lower energy-loss can also be potentially used for elemental mapping (Ahn & Krivanek, 1983). Figures 2a and 2b, show the raw spectra of the high-loss M_4,5_ edge region (onset at 883 eV) and the intermediate-loss N_4,5_ edge region (onset at 110 eV) for cerium, respectively. The N_4,5_ edge spectrum was acquired at a fraction of the dose of the M_4,5_ edge spectrum. The two distinct white lines of the M_4,5_ edge prominently stand out from the background in figure 2a, in contrast to the N_4,5_ edge which is barely discernible from the background as a small bump in figure 2c (see where the arrow points). In figure 2d, after the background subtraction, the saw tooth shape of the N_4,5_ edge is apparently evident.

**Figure 2.**
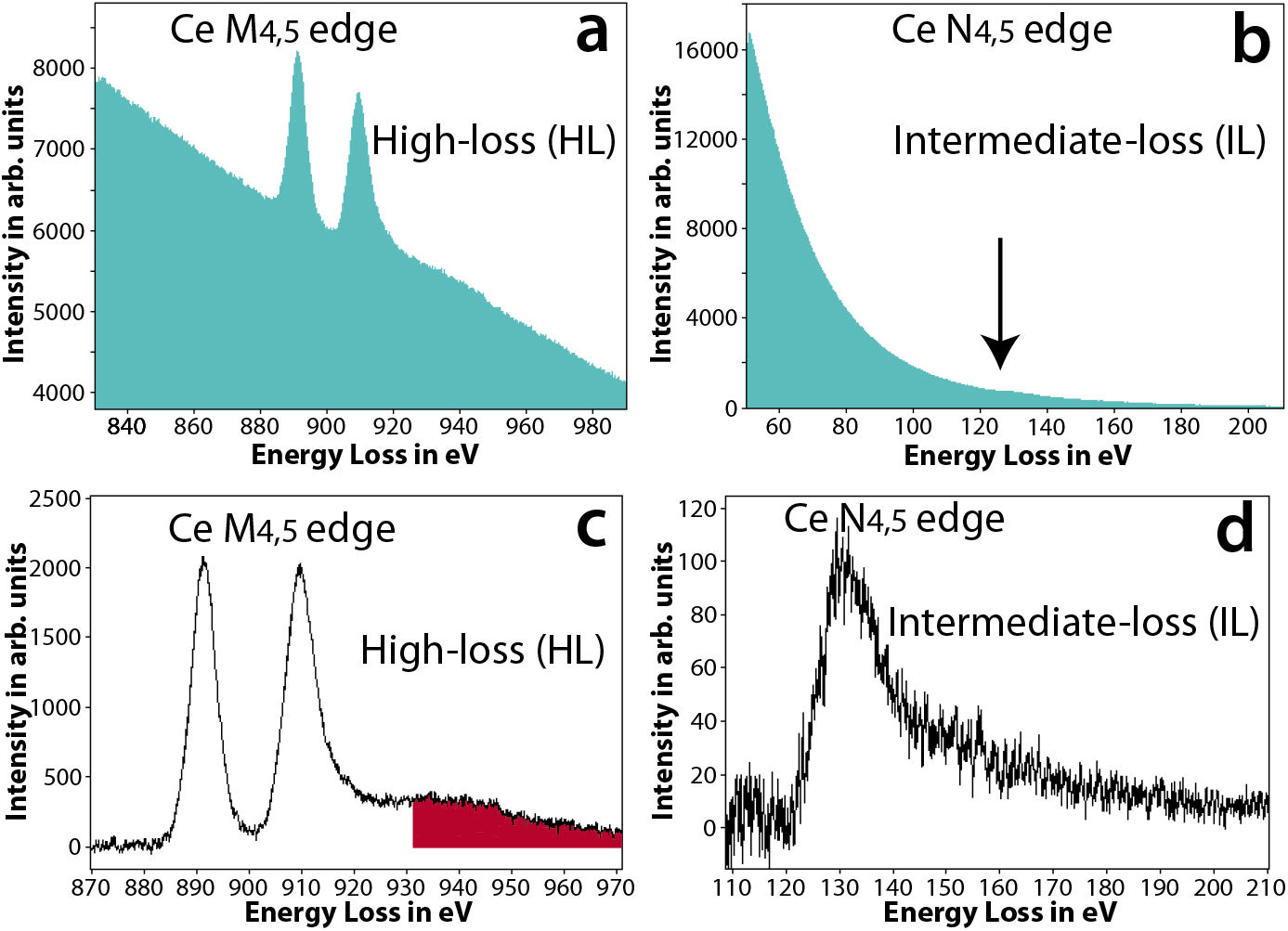
**a)** The raw spectrum acquired at the high-loss M_4,5_ edge of Ce, the two white line peaks with onset ~ 883 eV is clearly visible. **b)** The raw spectrum acquired at the intermediate-loss N_4,5_ edge of Ce, edge is barely visible as a small bump in the spectrum (see arrow direction). **c)** The background subtracted spectrum of (a), the M_4,5_ edge of Ce has a very high signal-above-background ratio (SBR). The red shaded region shows the Ce extended energy-loss fine structure (EXELFS) bleed into the Pr edge. **d)** The background subtracted spectrum of (b), the N_4,5_ edge of Ce is now clearly visible, though with a much lower SBR.

As outlined above, the total intensity and the SNR are significantly better at the intermediate-loss region than at the high-loss region. However, for edge visibility and detection, in addition to a good SNR, it is important to have a good signal-above-background ratio (SBR) (Egerton, 1996). Therefore, if the concentration of the lanthanide metal is low, then in spite of the good SNR, the SBR of the N_4,5_ edge can be so low that it cannot be distinguished from the background, and obtaining an elemental map at this edge will not be feasible. Also, it should be emphasized that, unlike M_4,5_ which is distinct and well separated for the lanthanides, the N_4,5_ edge have significant overlap from edge of adjacent lanthanides. The instance, the N_4,5_ edge of Ce (onset at 110 eV) will be difficult to distinguish from N_4,5_ edge of Pr (onset at 113 eV), for a more detailed discussion on this please see the supplementary section ‘Criteria for edge selection in EFTEM elemental mapping’. However, this should not limit one to obtain EFTEM elemental maps at of lanthanides at N_4,5_ edge, when there is no overlapping edge from other lanthanides or endogenous elements in the sample.

Figure 3a shows the conventional TEM image of mitochondrial matrix-APEX2 labeled with Ce2-DAB (second generation), figures 3b and 3b’ show the Ce spectra of high-loss M_4,5_ edge and the intermediate-loss N_4,5_ edge obtained on this region. A high-loss and an intermediate-loss elemental map were acquired on the sample, at the same dose rate and total dose (total dose for the high-loss acquisition was ~3% higher than the intermediate-loss acquisition). Figures 3c - c” and 3d – d’’ show the aligned and summed pre-edges and post-edge of the high-loss and intermediate-loss acquisition, respectively. It is very clear comparing these two sets of images that the pre-edges and the post-edge of the intermediate-loss show superior SNR and detail compared to the high-loss, as is expected. For the intermediate loss, the mitochondria in the post-edge image (figure 3d’’) looks sharper than the pre-edge images (figure 3d and d’) because the mitochondrial matrix is loaded with Ce, and there is a increase in signal in this region relative to the background. However, it is for the high-loss, that the difference between the post-edge image (Figure 3c”) and the pre-edge images (Figure 3c and 3c’) are really striking. The pre-edge images at high-loss are very blurry and noisy in comparison to the post-edge. The reason for this striking difference at the high-loss, is because of the much higher SBR of the M_4,5_ edge in comparison to the N_4,5_ edge. The SBR measured from the spectra for the signal and background integrated for a 30 eV window, is 1.0 and 0.3 for the high-loss and the intermediate-loss region, respectively.

**Figure 3.**
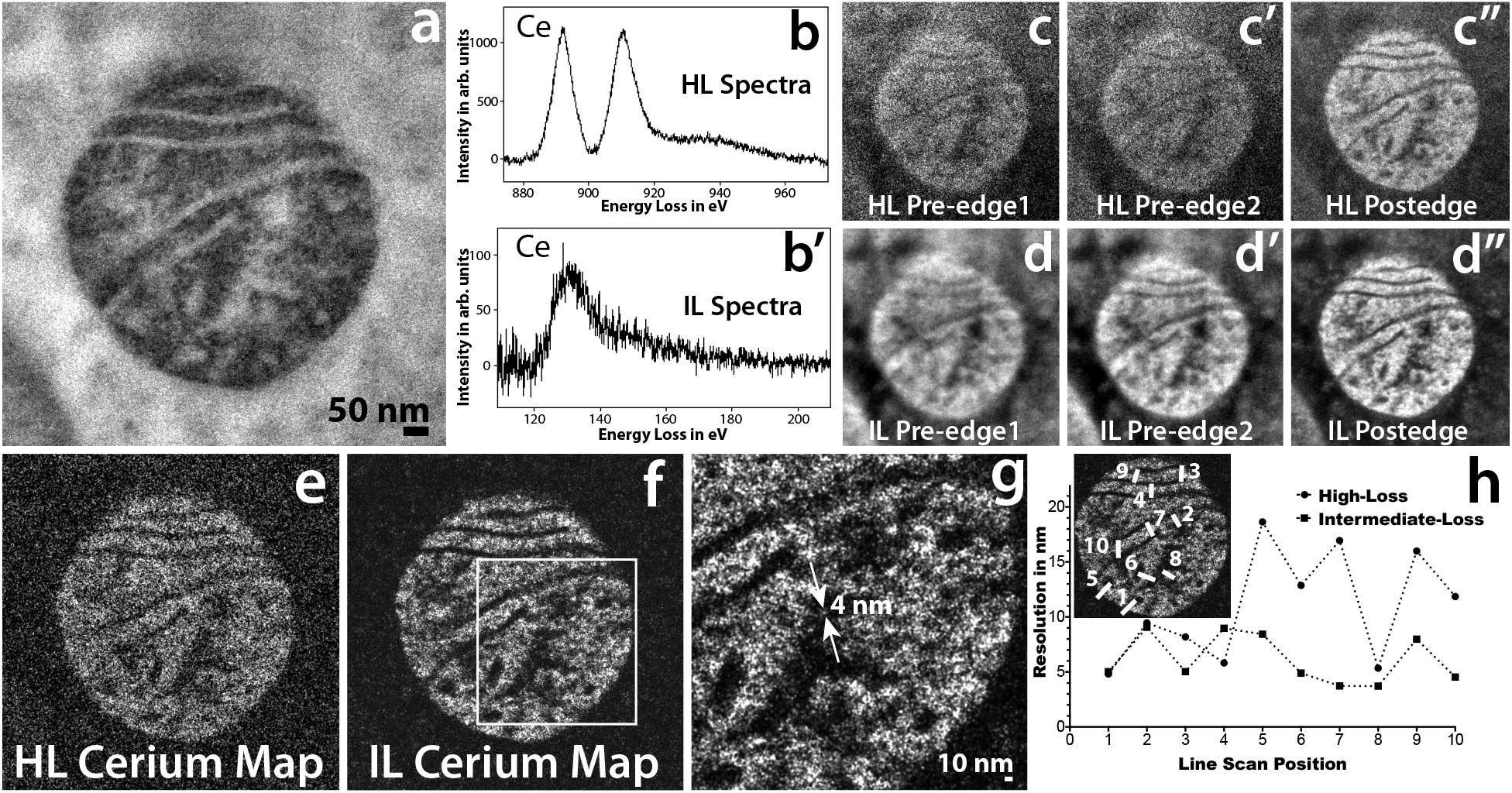
**a)** Conventional TEM image of mitochondrial matrix-APEX2 labeled with Ce2-DAB (second generation). **b)** The background subtracted high-loss spectrum acquired on the sample showing the M_4,5_ edge of Ce. **b’)** The background subtracted intermediate-loss spectrum acquired on the sample showing the N_4,5_ edge of Ce. **c, c’ and c’’)** The pre-edge1, pre-edge 2 and post-edge acquired for the high-loss M_4,5_ cerium edge at 815 eV, 855 eV and 899 eV respectively. Each energy-plane is a sum of 3 images acquired at an exposure of 100 sec/image. **d, d’ and d’’)** The pre-edge1, pre-edge 2 and post-edge acquired for the intermediate-loss N_4,5_ cerium edge at 75 eV, 98 eV and 138 eV respectively. Each energy-plane is a sum of 29 images acquired at an exposure of 10 sec/image. The dose and dose rate was nearly the same for both the high-loss and intermediate-loss acquisitions. **e)** The High-loss Ce elemental map. **f)** The intermediateloss Ce elemental map. **g**) Magnified view of the intermediate-loss Ce elemental map. **h)** The relative resolution comparison between the high-loss and the intermediate-loss Ce elemental map, measured by as the width of the intensity between 25 to 75% of the peak, for 10 different regions taken exactly the same position of the sample for both the maps (see the inset).

Figure 3e and 3f, show the elemental maps computed by the 3-window method for the high-loss and the intermediate-loss, respectively. Both, of which were acquired as a series with shorter exposure/energy plane. The intermediate-loss was acquired at a exposure of 10 sec/image, the high-loss required much longer exposures of 100 sec/image to be able to accumulate enough signal, the complete details of the acquisition can be found in the supplementary section. The intermediate-loss elemental map is clearly a lot sharper with a higher SNR, the SNR for the intermediate-loss map was 7.4 and for the high-loss map was 1.8, when calculated on the same region of the respective maps. It has been suggested, that in a plot of the intensity distribution across an edge in a image, the measure of the width of the intensity between 0.25 to 0.75 of its peak value, can be appropriated as the resolution (Reimer & Kohl, 2008). The edge width being proportional to the beam broadening, due to the point spread function (PSF) of the imaging system. The region between the matrix and the cristae or the exterior background in the elemental maps (figures 3e and 3f), serve as an edge for the measurement of the achieved resolution in the intermediate-loss and high-loss map. It should be cautioned, that unlike grayscale conventional TEM/STEM images, where the intensity on either side of the edge is fairly uniform, the pixel intensity of elemental maps vary significantly from one pixel to another because the intensity is proportional to the concentration of the element in that pixel. Therefore, measuring resolution from the edge step profile for elemental maps can be misleading. However, if the edge step profile is taken on exactly the same region of the image for both the high-loss and the intermediate-loss maps, then this can be used as a measure of relative degradation of resolution between the two maps. We obtained edge profiles at 10 exactly same locations of the sample on both the maps, width (nm) of the intensity between 0.25 to 0.75 of its peak value, are shown in figure 3e.

The resolution from this calculation, for the high-loss and the low-loss maps are 11.0 ± 3.0 and 6.1 ± 2.0 nm, respectively. This is consistent with the magnified view of the intermediate-loss elemental map in Figure 3g, which clearly shows a particle of 4 nm (measured as FWHM) distinguishable from the background. We would like to iterate, that this only implies that the resolution of the map acquired at the intermediate-loss region is ~ 1.8 times better than the map acquired at the high-loss region, and do not make any claim that these values are the absolute achievable resolution for the maps acquired at the respective energy-loss region.

As demonstrated above, both the SNR and the resolution of the elemental map is somewhat better at intermediate-loss than at high-loss. The correctness of the power law background subtraction at the intermediate-loss is not well known. However, there can be some speculation that the contrast that is seen in the intermediate-loss elemental map may actually be just density contrast of the heavily stained region containing cerium + ruthenium (secondary fixative), erroneously being interpreted as a cerium elemental map. On the other hand, for the high-loss 3-window method (energy-loss > 200 eV), the accuracy of the power-law background subtraction, and the optimization of the position and width of the energy-loss EFTEM images are well established (Hofer, et al., 1997). Therefore, there needs to be additional validation that the contrast that is seen in the intermediate-loss Ce map, is indeed an elemental contrast. To test this hypothesis, a control sample was made, by labeling mitochondrial matrix-APEX2 with plain DAB. The intensity of the Mitomatrix staining with DAB was equally like the Ce2DAB sample, but contains only ruthenium and no cerium. If an intermediate-loss map is acquired on the control sample with the same experimental settings as that of the mitochondrial matrix-APEX2 labeled with Ce2-DAB (figure 3f), and if the map shows positive contrast for the matrix region of the mitochondria, then it can be concluded that contrast seen in figure 3f is not a true Ce elemental contrast. Figure 4a - d, shows a high-loss and intermediate-loss map acquired using the 3-window method on the control sample, with the same spectrometer settings as in figure 3 (see the supplementary section for details). The mitochondrial matrix in he high-loss map (figure 4c) shows a faint positive contrast indicating an under subtraction of the background, and in the intermediate-loss map (figure 4d), it shows a negative contrast due to over subtraction of the background. Also, in figure 4d, although the mitochondrial matrix shows negative contrast, the ribosome show positive contrast (see where the arrows point). Therefore, it can be concluded that in this case, mitochondrial matrix showing a negative contrast and not a positive contrast (i.e. density contrast), would have led to a partial suppression of the Ce elemental signal, if the sample did contain Ce (control sample does not contain Ce). Figures 4e - h, illustrates another example of high-loss and intermediate-loss elemental maps acquired on the control sample by the 3-window method, in this case the intermediate-loss map was acquired at a slightly different spectrometer settings than the previous example in figure 4d. The intermediate-loss map was acquired for slit width of 15 eV instead of 20 eV, and the pre-edge 1, pre-edge 2 and post-edge were acquired at 84 eV, 102 eV and 135 eV, respectively. In the previous example (figure 4d), the pre-edge 1, pre-edge 2 and post-edge were acquired at 75 eV, 98 eV and 138 eV, respectively. The high-loss map (figure 4g) shows a faint positive contrast as in the previous example (figure 4c), however, the intermediate-loss map (figure 4h) now shows a positive contrast instead of a negative contrast for the mitochondria. It can be deduced from figures 4d and 4h, that the intermediate-loss elemental maps are more susceptible to inconsistencies in background subtraction even for small changes in the slit width or position. The lower SBR of the intermediate-loss N_4,5_ edge, is the primary reason for this contrast inversion.

**Figure 4.**
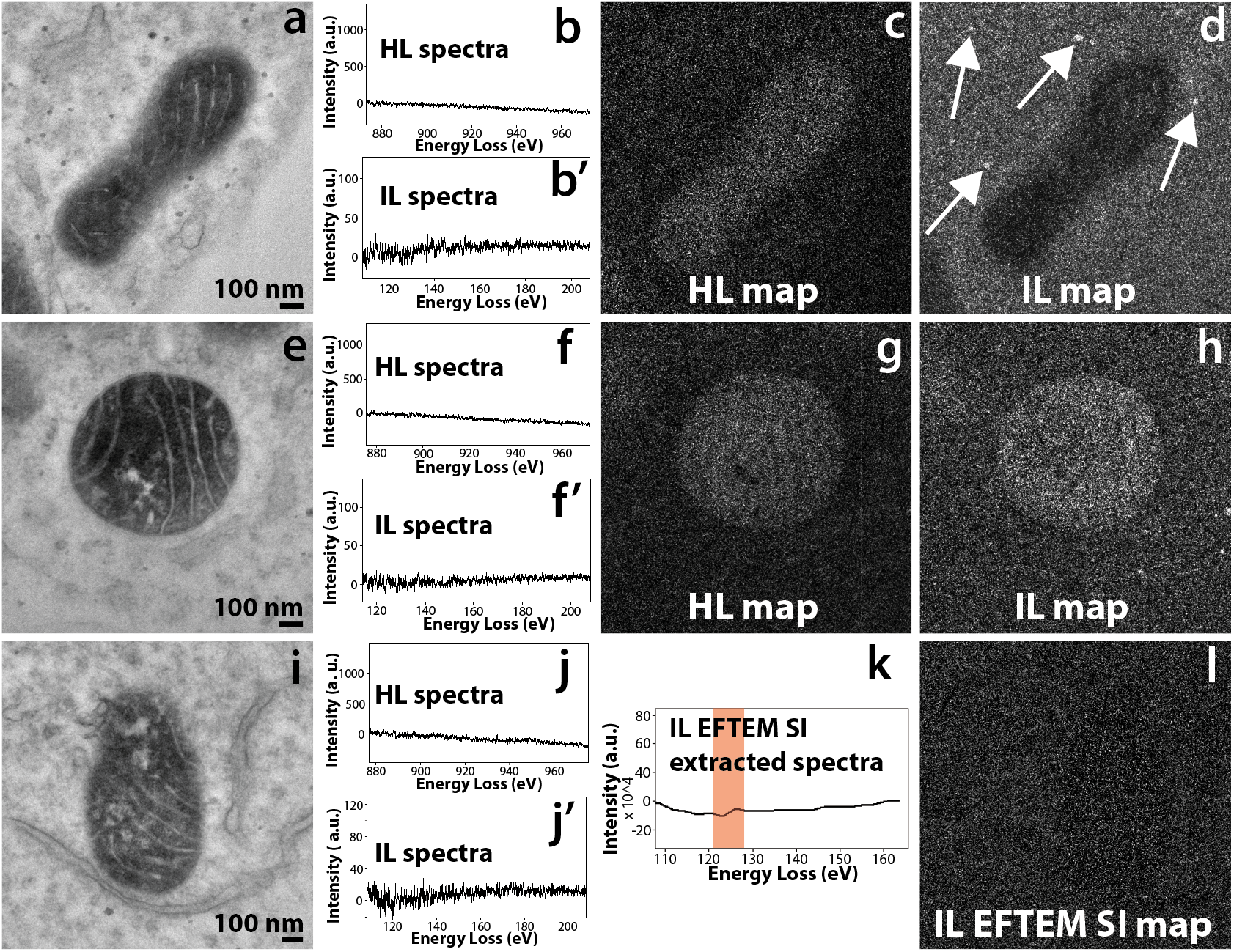
**a)** Conventional TEM image of the control sample of mitochondrial matrix-APEX2 labeled with plain DAB (no Ce or any other lanthanides). **b and b’)** The background subtracted high-loss and intermediate-loss spectrum acquired on the mitochondrial matrix region of (a), confirming the absence of Ce in the sample. **c)** The high-loss elemental map computed by the 3-window method, for pre edge 1, pre edge 2 and post-edge obtained for a slit width of 30 eV at 815 eV, 855 eV and 899 eV respectively. The image shows a positive contrast for the matrix region, implying an under subtraction of the background. **d)** The intermediate-loss elemental map computed by the 3-window method, for pre edge 1, pre edge 2 and post-edge obtained for a slit width of 20 eV at 75 eV, 98 eV and 138 eV respectively. The image shows a negative contrast for the matrix region, implying an over subtraction of the background. However the ribosome show a positive contrast (see the arrows), implying inconsistencies in background subtraction. **e)** Conventional TEM image of a different region of the same control sample. **f and f)** The background subtracted high-loss and intermediate-loss spectrum acquired on the mitochondrial matrix region of (d), confirming the absence of Ce in the sample. **g)** The high-loss elemental map computed by the 3-window method, for pre edge 1, pre edge 2 and post-edge obtained for a slit width of 30 eV at 815 eV, 855 eV and 899 eV respectively. The image shows a positive contrast for the matrix region, implying an under subtraction of the background. **h)** The intermediate-loss elemental map computed by the 3-window method, for pre edge 1, pre edge 2 and post-edge obtained for a slit width of 15 eV at 84 eV, 102 eV and 135 eV respectively. The image shows a positive contrast for the matrix region, implying an under subtraction of the background. **i)** Conventional TEM image of a different region of the same control sample. **j and j’)** The background subtracted high-loss and intermediate-loss spectrum acquired on the mitochondrial matrix region of (g), confirming the absence of Ce in the sample. **k)** The virtual intermediate-loss spectrum extracted from the EFTEM SI image-series, corroborating the results in (h’). **l)** The intermediate-loss elemental map computed by the EFTEM SI method, for an acquisition acquired from 90 eV to 150 eV, for a slit width of 8 eV and an energy step of 3 eV. The signal was integrated from the region represented by the red box in k. The image shows a no contrast and only noise for the matrix region, implying a correct subtraction of the background.

From the discussion presented above, it is evident that the best technique to acquire an elemental map at the intermediate-loss region is the EFTEM Spectrum Imaging (EFTEM SI) (Schaffer, et al., 2008; Schaffer, et al., 2006; Terada, et al., 2001; Watanabe & Allen, 2012). This technique involves acquiring a series of EFTEM images with a narrow slit, and with the spectrometer continuously stepping through energies (or energy-loss), from a region representing the edge to the background region prior to the edge onset. The spectrometer is stepped in such a way, that there is a slight energy overlap between successive acquisitions. From the thus acquired EFTEM SI image series, a spectrum can be virtually extracted using the SI picker tool in Digital Micrograph (Gatan, Inc). Since this extracted spectrum directly correlates to the image series, the user has complete flexibility to choose which region (i.e. which subset of images) contributes to the background and which region contributes to the elemental signal. Additionally, the user can compensate for any chemical shifts or spectrum energy shifts post acquisition.

Figure 4i – l, shows the EFTEM SI intermediate-loss acquisition on the same control sample shown in the previous two examples (figures 4a – h). Figures 4j and 4j’, is the high-loss and intermediate-loss background subtracted spectra, that confirm that there is no Ce in the sample. Figure 4k, is the background subtracted virtual spectrum extracted from the EFTEM SI image series, also confirming absence of Ce. Figure 4l, displays the Ce intermediate-loss elemental map that was computed from the energy-loss region indicated by the red box in Figure 4k, i.e. the region that would have contained the N_4,5_ edge of Ce (the control does not contain Ce). This map shows neither a positive or a negative contrast at the mitochondrial matrix region and is only noise, clearly authenticating the accuracy of background extrapolation of the EFTEM SI technique over the 3-window method.

Figure 5, illustrates some examples of EFTEM SI intermediate-loss elemental maps of three different cellular targets, namely using Mitomatrix-APEX2, histone H2B-Nucleosome and EdU-DNA labeled with three different lanthanides: cerium, lanthanum and neodymium, respectively.

**Figure 5.**
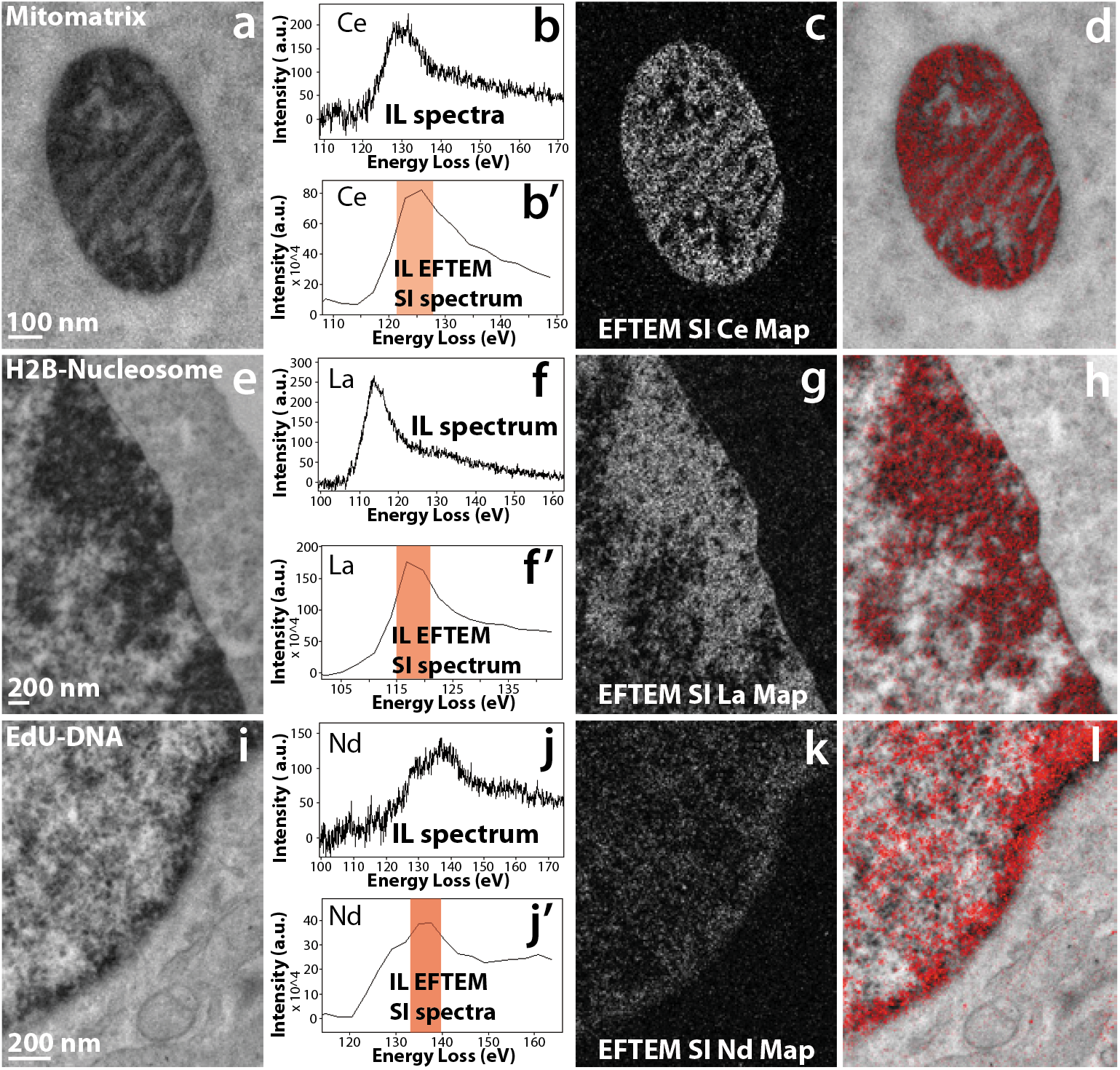
**a)** Conventional TEM image of mitochondrial matrix-APEX2 labeled with Ce2-DAB (second generation). **b)** The background subtracted intermediate-loss spectrum acquired on the mitochondrial matrix region of (a), confirming the presence of Ce in the sample. **b’)** The virtual intermediate-loss spectrum extracted from the EFTEM SI imageseries, corroborating the results in (b). **c)** The intermediate-loss Ce elemental map computed by the EFTEM SI, the region (or images) contributing to the signal is enclosed in the red box in (b’). **d)** Single Color EM. **e)** Conventional TEM image of MiniSOG-H2B labeled with La2-DAB (second generation). **f)** The background subtracted intermediate-loss spectrum acquired on the nucleus of (d), confirming the presence of La in the sample. f) The virtual intermediate-loss spectrum extracted from the EFTEM SI image-series, corroborating the results in (e). **g)** The intermediate-loss La elemental map computed by the EFTEM SI, the region (or images) contributing to the signal is enclosed in the red box in (f). **h)** Single Color EM. **i)** Conventional TEM image of DNA incubated with EdU and clicked with Fe-TAML-azide for oxidation of Nd-DAB2 (first generation lanthanide DAB). **j)** The background subtracted intermediate-loss spectrum acquired on the nucleus of (g), confirming the presence of Nd in the sample. **j’)** The virtual intermediate-loss spectrum extracted from the EFTEM SI image-series, corroborating the results in (h). **k)** The intermediate-loss Nd elemental map computed by the EFTEM SI, the region (or images) contributing to the signal is enclosed in the red box in (j’). **l)** Single Color EM.

Figure 5 a – d, demonstrates the acquisition of EFTEM SI intermediate-loss cerium elemental map on the mitochondrial matrix-APEX2 labeled with Ce2-DAB. Figure 5b shows the real background subtracted intermediate-loss spectrum acquired on the mitochondrial region of (a), showing presence of Ce. In Figure 5b’, the virtual spectrum extracted from the EFTEM SI image-series, and subsequently background subtracted can be seen. Though the extracted spectrum of 5b’ is of a lower energy resolution than the actual spectra of 5b, the two match very well on the general shape and profile of the N_4,5_ edge. Computing the elemental map with the aid of the spectrum in b’, with the flexibility to choose parameters post-acquisition, gives undisputable confidence in the correctness of the final result. Figure 5c shows the Ce elemental map, computed from the energy-loss region indicated by the red box and figure 5d displays the Color EM image of the pseudocolored elemental map overlaid over the conventional TEM image. The Color EM image was generated by an ImageJ plugin that we developed, and this algorithm has been described elsewhere (Adams, et al., 2016).

The main drawback of acquiring elemental maps by the EFTEM SI method over the 3-window method, is that it necessitates the use a narrow slit width and with an energy overlap between successive images, thereby requiring far more electron dose. For e.g., the EFTEM SI intermediate-loss elemental map acquisition in figure 5c, required 2.6 times the dose of the 3-window acquisition in figure 3f, for an elemental map with much poorer SNR. The SNR of the EFTEM SI elemental map in figure 5c is 1.6 in comparison to the SNR of 7.4 for the 3-window method in figure 3f. However, the EFTEM SI acquisition in this case was oversampled, and considerable reduction in dose can be achieved by making very small changes in the acquisition parameters. To illustrate, the EFTEM SI intermediate-loss elemental map in figure 5c, was acquired between a energy-loss of 90 eV to 150 eV, with a slit width of 8 eV and an energy step of 3 eV, resulting in a total of 21 images. The region that used for background subtraction was 98.6 eV to 107.1 eV and that for the Ce signal was 121.4 eV to 127.1 eV. The regions from 90 to 98.5 eV and 121.2 to 150 eV did not contribute in any way for the computation of the elemental map. Therefore, an acquisition from ~ 94 eV to 134 eV, should be sufficient and is wide enough to compensate for any chemical shifts (−2 eV, +7 eV). An EFTEM SI acquisition with a 5 eV energy step instead of 3 eV, would reduce the total number of images in the EFTEM SI stack from 21 to 9, reducing the total dose by ~ 60%. This would then make dose required for the EFTEM SI acquisition similar to the dose required for the 3-window method.

Figures 5e – 5f, illustrates the EFTEM SI intermediate-loss lanthanum elemental map acquisition on the MiniSOG-H2B labeled with La2-DAB. The actual spectrum (figure 5f) and the spectrum extracted from the EFTEM SI stack (figure 5f) show very good correlation in shape and profile. The lanthanum elemental map shows good signal with a clean background in figure 5g, and figure 5h shows the corresponding color overlay.

Figures 5i – 5l, shows the EFTEM SI intermediate-loss neodymium elemental map acquired on the EdU-DNA labeled with Nd-DAB2. The actual spectrum and the extracted EFTEM SI spectrum show good correlation similar to the previous two examples, but the signal in the neodymium elemental map (figure 5k) is faint. The reason for the poor neodymium signal is that, unlike the previous two examples, this sample was labeled with first generation lanthanide DAB rather than the second generation. Therefore, in this case there is 4 times lesser metal (i.e. Nd) per DAB, due to which there is a much lower signal amplification, the signal-above-background (SBR) is very low. To accommodate for the low SBR, 3 sets of EFTEM SI stack were acquired on the sample, subsequently aligned and added to create the summed EFTEM SI stack to improve the SNR (Watanabe & Allen, 2012). Therefore, even for cases where the concentration of the lanthanide is low, EFTEM SI method of acquiring intermediate-loss elemental maps is still possible.

## CONCLUSION

The method for multi-color or single-color EM that we previously developed, was achieved by the sequential deposition of specific lanthanide chelates conjugated to diaminobenzidine, which were selectively oxidized by orthogonal photosensitizers / peroxidases. The Color EM was generated by the pseudo-colored overlay of the core-loss or high-loss EFTEM elemental map of the lanthanide onto the conventional image. The lanthanides, in addition to the high-loss M_4,5_ edge, also exhibit an intermediate-loss N_4,5_ edge that has significantly higher signal but suffers from very low signal-above-background (SBR), which generally limits the acquisition of elemental maps at the intermediate-loss. The synthesis of the new second generation lanthanide DABs, which contains 4 times more lanthanide per DAB, gives significant signal amplification and improvement to the SBR, making collection of elemental maps at the intermediate-loss region more feasible. For the same experimental conditions, the intermediate-loss provides ~ 4x improvement in SNR and ~ 2x improvement in resolution in comparison with the high-loss region. The 3-window method of computing elemental maps is prone to errors due to inconsistencies in background subtraction. The EFTEM SI method of generating elemental maps, provides results with very high signal fidelity and confidence, however at the expense of SNR. Also, Multi color EM of mapping 2 or more lanthanides is difficult at the intermediate-loss region because of significant overlap of edges between adjacent lanthanides. However, if there is sufficient Z gap between the lanthanides, for e.g. lanthanum and gadolinium or cerium and ytterbium, then multi Color EM should be potentially possible in the intermediate-loss region and will be explored in the future.

## ACKNOWLEDGEMENTS

We would like to thank Rob Bilhorn and Benjamin Bammes of Direct Electron for help with the DE-12 detector. We would like to thank David Mastronarde of University of Colorado Boulder for help with SerialEM. This work was supported by NIH grants R01GM086197 and R24GM137200.

## SUPPLEMENTARY INFORMATION

**Supplementary Figure 1:**
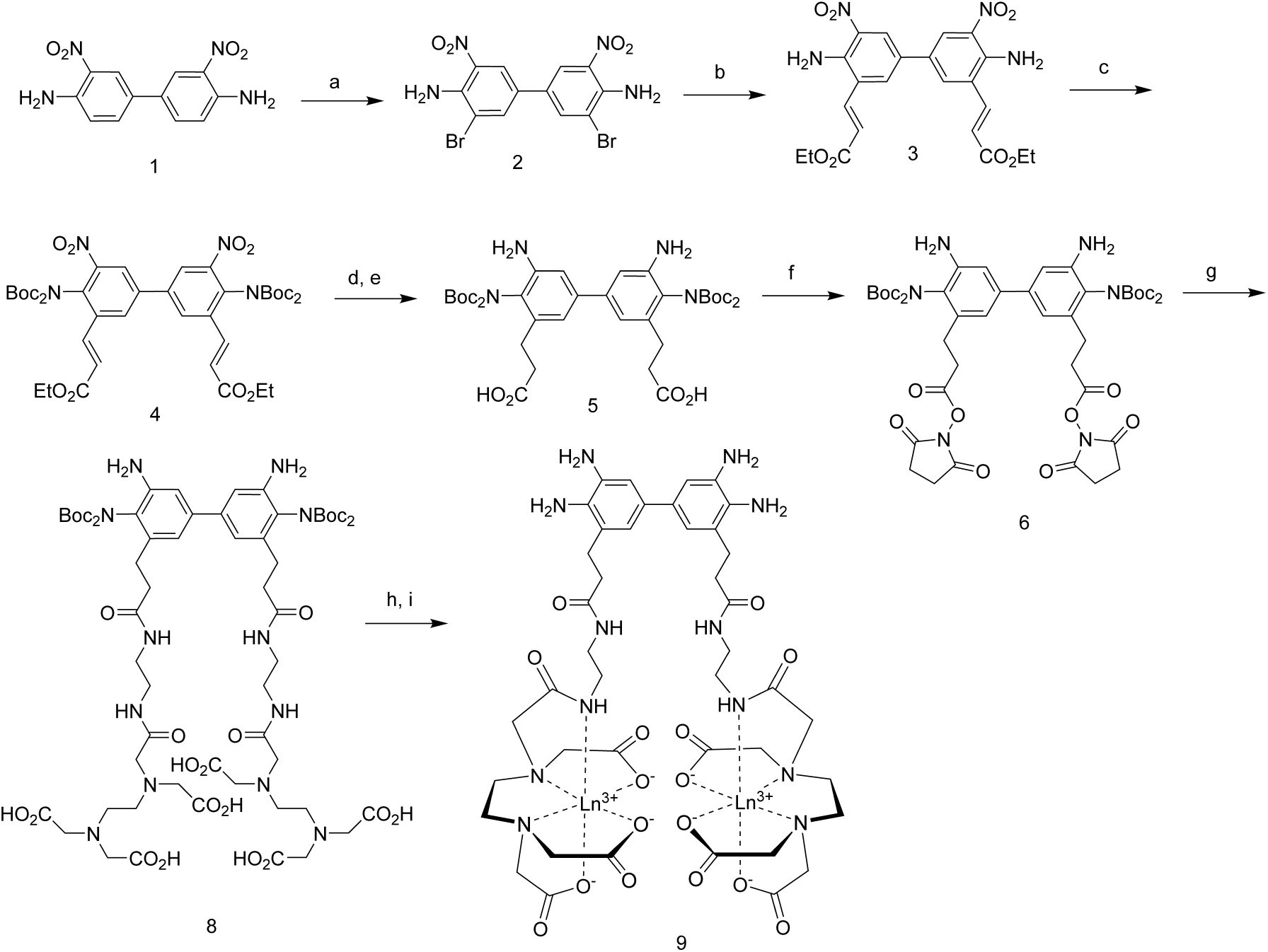
Scheme of the synthesis of Ln2-DAB. Reagents, solvents, conditions, a) Br_2_, acetic acid, rt; b) ethyl acrylate, Pd(OAc)_2_, (o-tolyl)_3_phosphine, NEt_3_, DMF, 80°C; c) BOC anhydride, DMAP, DMF, 80°C; d) 1 atm H_2_, Pd-C, EtOAc-EtOH, rt; e) NaOH, dioxane-MeOH, rt; f) HOSu, EDC, DMF, rt; g) eda-EDTA, NEt_3_, DMSO, rt; h) TFA, rt; i) LnCl_3_, pH 7 buffer, rt.

### Criteria for edge selection in EFTEM elemental mapping

An element can have several ionization edges, some being major edges and others being minor edges. For example, cerium (Ce) has major edges at onset of 20 eV (O_2,3_), 110 eV (N_4,5_), 883 eV (M_5_) and 901 eV (M_4_); minor edges at onset of 207 eV (N_2,3_), 290 eV (N_1_), 1185 eV (M_3_), 1273 eV (M_2_) and 1435 eV (M_1_) (Ahn & Krivanek, 1983). The edge onset energy can vary by about (−2 eV, +7 eV) due to bonding effects called chemical shifts, in addition to the spread due to chromatic aberration of the imaging system, which can range from few eV to tens of meV depending on the instrument (Carter & Williams, 2009). The edge that is selected for elemental mapping/quantification needs to fulfill certain requirements. Firstly, the edge should be a major edge in the range ~ 100 - 2000 eV (Egerton, 1996), the optimum EELS energy region being within 1000 eV (Ahn & Krivanek, 1983). Secondly, the edge should have a distinct shape, for e.g. a sawtooth or peaked at threshold like white-line for easy identification (Egerton, 1996). Finally, the edge location (in eV) should be distinct with no overlapping edge due to other adjacent elements in the periodic table.

For Ce, as described above there are potentially 4 major edges, however the first major edge O_2,3_ at 20 eV does not satisfy any of the three conditions. However, using complex instrumentation like monochromators fitted to a state of the art TEM and with the aid of mathematical tools such as nonlinear multivariate statistical analysis, ability to differentiate material compositions based on subtle differences in edge < 50 eV has been shown (Pfannmoller, et al., 2012; Pfannmoller, et al., 2011).

The N_4,5_ edge (see figure 2b and 2d) satisfies the first two conditions, it is a major peak > 100eV and has a sharp and distinct sawtooth shape. However, the N_4,5_ edge for Ce is at 110 eV, and for praseodymium (Pr) it is at 113 eV, and due to the chemical shifts and the chromatic aberrations of the imaging system, unambiguous identification of the element solely based on this edge is not possible. The last two of these edges, the M_5_ and the M_4_ edge are very close to each other (see figure 2a and 2c), and for the purpose of elemental mapping, they are generally considered together as the M_4,5_ edge. The M_4,5_ edge for Ce has the so called white lines shape, that starts at ~ 883 eV and extends to ~ 920 eV. This edge is sufficiently separately from the M_4,5_ edge of Pr, which starts only at ~931 eV is the preferred choice for elemental mapping/quantification. in fact, we have previously shown a multicolor EM of two astrocyte processes contacting the same synapse, with one astrocyte being labeled with Cerium conjugated DAB and the other with Praseodymium conjugated DAB (Adams, et al., 2016). It should be pointed out that, although the Cerium M_4,5_ edge is explicitly separated from the Praseodymium, the extended energy-loss fine structure (EXELFS) of Ce bleeds into the Pr edge and beyond (see the area shaded in red in figure 2c), this spectral bleed-through has to be mathematically subtracted when maps of adjacent elements in the periodic table are computed (Adams, et al., 2016).

Alternatively, if there is already a priori information on the elemental composition of the sample, and the primary purpose of using EELS/EFTEM is not element identification but localization or distribution of a particular element in the sample, then the criteria for the edge selection for the particular element can be slightly relaxed. For e.g., if a single cellular protein/organelle in cells was labeled by the deposition of only one lanthanide conjugated to diaminobenzidine, and the localization of this protein/organelle is to be visualized by acquiring an EFTEM elemental map of the specific lanthanide metal, in a so called Single color EM. In such a scenario, the elemental map for the lanthanide can be potentially acquired on the intermediate-loss region of the N_4,5_ edge instead of the high-loss region of the M_4,5_ edge, provided that there are no overlapping edge from any of the endogenous elements in the sample.

### EFTEM acquisition parameters

For the high-loss EFTEM images of the mitochondrial matrix-APEX2 labeled with Ce2-DAB by the 3-window method (See figure 3e), a series of 3 images with a 100 s exposure/image, was acquired by the DE-12 detector at a frame rate of 0.04 fps and a magnification of 10kX (pixel size 0.5 nm) for each energy window. The pre-edge1, pre-edge2 and post-edge were acquired for a slit width of 30 eV at energy shifts of 815, 855 and 899 eV, respectively. The EFTEM images were acquired at a dose rate of ~ 1.2 × 10^-4^ PA/nm^2^ and the total dose for the acquisition was ~ 6.8 × 10^5^ e^-^/nm^2^.

For the intermediate-loss EFTEM images of the mitochondrial matrix-APEX2 labeled with Ce2-DAB by the 3-window method (See figure 3f), a series of 29 images with a 10 s exposure/image, was acquired by the DE-12 detector at a frame rate of 2 fps and a magnification of 10kX (pixel size 0.5 nm) for each energy window. The pre-edge1, pre-edge2 and post-edge were acquired for a slit width of 20 eV at energy shifts of 75, 98 and 138 eV, respectively. The EFTEM images were acquired at a dose rate of ~ 1.2 × 10^-4^ PA/nm^2^ and the total dose for the acquisition was ~ 6.6 × 10^5^ e^-^/nm^2^.

For the high-loss EFTEM images of the control sample of mitochondrial matrix-APEX2 labeled with plain DAB by the 3-window method (See figure 4c), a series of 5 images with a 100 s exposure/image, was acquired by the DE-12 detector at a frame rate of 0.04 fps and a magnification of 10kX (pixel size 0.5 nm) for each energy window. The pre-edge1, pre-edge2 and post-edge were acquired for a slit width of 30 eV at energy shifts of 815, 855 and 899 eV, respectively. The EFTEM images were acquired at a dose rate of ~ 1.2 × 10^-4^ PA/nm^2^ and the total dose for the acquisition was ~ 1.1 × 10^6^ e^-^/nm^2^.

For the intermediate-loss EFTEM images of the control sample of mitochondrial matrix-APEX2 labeled with plain DAB by the 3-window method (See figure 4d), a series of 30 images with a 10 s exposure/image, was acquired by the DE-12 detector at a frame rate of 2 fps and a magnification of 10kX (pixel size 0.5 nm) for each energy window. The pre-edge1, pre-edge2 and post-edge were acquired for a slit width of 20 eV at energy shifts of 75, 98 and 138 eV, respectively. The EFTEM images were acquired at a dose rate of ~ 1.2 × 10^-4^ PA/nm^2^ and the total dose for the acquisition was ~ 6.8 × 10^5^ e^-^/nm^2^.

For the high-loss EFTEM images of the control sample of mitochondrial matrix-APEX2 labeled with plain DAB by the 3-window method (See figure 4g), a series of 5 images with a 100 s exposure/image, was acquired by the DE-12 detector at a frame rate of 0.04 fps and a magnification of 10kX (pixel size 0.5 nm) for each energy window. The pre-edge1, pre-edge2 and post-edge were acquired for a slit width of 30 eV at energy shifts of 815, 855 and 899 eV, respectively. The EFTEM images were acquired at a dose rate of ~ 1.2 × 10^-4^ PA/nm^2^ and the total dose for the acquisition was ~ 1.1 × 10^6^ e^-^/nm^2^.

For the intermediate-loss EFTEM images of the control sample of mitochondrial matrix-APEX2 labeled with plain DAB by the 3-window method (See figure 4h), a series of 30 images with a 10 s exposure/image, was acquired by the DE-12 detector at a frame rate of 2 fps and a magnification of 10kX (pixel size 0.5 nm) for each energy window. The pre-edge1, pre-edge2 and post-edge were acquired for a slit width of 15 eV at energy shifts of 84, 102 and 135 eV, respectively. The EFTEM images were acquired at a dose rate of ~ 1.2 × 10^-4^ PA/nm^2^ and the total dose for the acquisition was ~ 6.8 × 10^5^ e^-^/nm^2^.

For the intermediate-loss EFTEM images of the control sample of mitochondrial matrix-APEX2 labeled with DAB by the 3-window method (See figure 4h), a series of 30 images with a 10 s exposure/image, was acquired by the DE-12 detector at a frame rate of 2 fps and a magnification of 10kX (pixel size 0.5 nm) for each energy window. The pre-edge1, pre-edge2 and post-edge were acquired for a slit width of 15 eV at energy shifts of 84, 102 and 135 eV, respectively. The EFTEM images were acquired at a dose rate of ~ 1.2 × 10^-4^ PA/nm^2^ and the total dose for the acquisition was ~ 6.8 × 10^5^ e^-^/nm^2^.

For the intermediate-loss EFTEM images of the control sample of mitochondrial matrix-APEX2 labeled with DAB by the EFTEM SI method (See figure 4l), the stack was acquired by the DE-12 detector from energy-loss of 165 eV to 87 eV, with a slit width of 8 eV and energy step of 3 eV. The stack was acquired at a magnification of 12 kX (pixel size 0.4 nm), frame rate of 0.1 fps and exposure of 60 s per individual energy plane. The EFTEM images were acquired at a dose rate of ~ 1.8 × 10^-4^ PA/nm^2^ and the total dose for the acquisition was ~ 1.9 × 10^6^ e^-^/nm^2^.

For the intermediate-loss EFTEM images of the mitochondrial matrix-APEX2 labeled with Ce2-DAB by the EFTEM SI method (See figure 5c), the stack was acquired by the DE-12 detector from energy-loss of 150 eV to 90 eV, with a slit width of 8 eV and energy step of 3 eV. The stack was acquired at a magnification of 12 kX (pixel size 0.4 nm), frame rate of 0.1 fps and exposure of 60 s per individual energy plane. The EFTEM images were acquired at a dose rate of ~ 1.8 × 10^-4^ PA/nm^2^ and the total dose for the acquisition was ~ 1.5 × 10^6^ e^-^/nm^2^.

For the intermediate-loss EFTEM images of the MiniSOG-H2B labeled with La2-DAB by the EFTEM SI method (See figure 5g), the stack was acquired by the DE-12 detector from energy-loss of 78 eV to 144 eV, with a slit width of 8 eV and energy step of 3 eV. The stack was acquired at a magnification of 10 kX (pixel size 0.5 nm), frame rate of 0.1 fps and exposure of 60 s per individual energy plane. The EFTEM images were acquired at a dose rate of ~ 2.6 × 10^-4^ PA/nm^2^ and the total dose for the acquisition was ~ 2.2 × 10^6^ e^-^/nm^2^.

For the intermediate-loss EFTEM images of the EdU-DNA labeled with Nd-DAB2 by the EFTEM SI method (See figure 5k), the stack was acquired by the DE-12 detector from energy-loss of 96 eV to 165 eV, with a slit width of 8 eV and energy step of 3 eV. The stack was acquired at a magnification of 10 kX (pixel size 0.5 nm), frame rate of 0.1 fps and exposure of 60 s per individual energy plane. The EFTEM images were acquired at a dose rate of ~ 2.6 × 10^-4^ PA/nm^2^ and the total dose for the acquisition was ~ 2.3 × 10^6^ e^-^/nm^2^.

